# ProtMiscuity: a database of promiscuous proteins

**DOI:** 10.1101/477380

**Authors:** Ana Julia Velez Rueda, Nicolas Palopoli, Matías Zacarías, Gustavo Parisi

**Affiliations:** Departamento de Ciencia y Tecnología, Universidad Nacional de Quilmes - CONICET, Roque Sáenz Peña 352, Bernal, B1876BXD, Buenos Aires, Argentina.

## Abstract

**Summary:** ProtMiscuity is a manually-curated database of promiscuous proteins. It is annotated with information about canonical and promiscuous activities comprising 88 different reactions in 57 proteins from 40 organisms. ProtMiscuity could assist in the study of the underlying mechanisms of promiscuous reactions by offering a collection of experimentally derived data, extensively linked with other databases providing biological, structural and functional information.

**Availability and Implementation:** The responsive web interface of ProtMiscuity provides support for easier navigation and visualization of the database contents on multiple devices. It is implemented in HTML, CSS, JavaScript, Angular4 and NodeJS. ProtMiscuity is hosted on our server and can be freely accessed at http://ufq.unq.edu.ar/protmiscuity

**Contact:** Gustavo Parisi

## Introduction

Even though protein promiscuity has been extensively studied in the last decades, the term itself is not well defined yet. It has been used to describe many different phenomena and different classification schemes has been proposed (Hult and Berglund, 2007; López-Iglesias and Gotor-Fernández, 2015). From a chemical and functional point of view, catalytic promiscuity may be described as the ability of an enzyme to catalyse a secondary activity at the same active site where its canonic or primary activity occurs. Accordingly, promiscuity can be classified by the chemistry of the reaction (Copley, 2003). Instead, Khersonsky and Tawfik (Khersonsky and Tawfik, 2010) described catalytic promiscuity as the capability of an enzyme to catalyze a different reaction than that for which the protein has evolved. Besides their many definitions and perspectives, promiscuity is not such an uncommon phenomenon as previously thought, and is increasingly permeating into drug discovery protocols, organic synthesis, pharmacology and biotechnology (Nobeli *et al.*, 2009).

In spite of its biological relevance and functional diversity, there is still no publicly available collection of scientific evidence on protein protmiscuity. Here we present ProtMiscuity, an online database that aims to fill this gap by providing a manually curated dataset of promiscuous enzymes and related information.

## Features

ProtMiscuity is a curated database of promiscuous proteins that aims to centralize experimentally characterized examples of this phenomenon. An initial dataset of relevant proteins and associated publications was developed through the implementation of web-scraping on PubMed (https://www.ncbi.nlm.nih.gov/pubmed/) and text-mining techniques over this bibliography, using standard libraries in the Python programming language. This collection of putative references to promiscuous proteins was inspected to filter out unconvincing cases by careful consideration of the available evidence, including data collected manually from other publications and databases. This process resulted in a total of 57 proteins with one or more characterized promiscuous activities. These proteins are described by their UniProt identifiers (Pundir *et al.*, 2017) and correspond to 2001 protein chain structures in the PDB (Touw *et al.*, 2015). Reactions, both promiscuous and canonical, are characterized in ProtMiscuity by information obtained from the literature regarding known substrates and products, *Km* and *kcat* values, active site residues and reaction conditions. Likewise, substrates and products related to each described reaction were linked to the information available in PDB Ligand Expo (Sitzmann *et al.*, 2012) and PubChem (Kim *et al.*, 2016) to facilitate the identification of possible ligands by chemical similarity. ProtMiscuity covers a total of 88 described chemical reactions in proteins coming from 40 different organisms. Among them, ~68% have only one promiscuous reaction, while 20% of the entries have two and ~6% have three or four promiscuous activities.

In order to provide the users with further structural and functional information, each protein is linked to resources such as the CoDNaS database of conformational diversity (Monzon *et al.*, 2016), KEGG pathways (Tanabe and Kanehisa, 2012), Catalytic Site Atlas annotations (Furnham *et al.*, 2014) and QuickGO terms (Binns *et al.*, 2009). ProtMiscuity also includes a tutorial section and answers to frequently asked questions to facilitate navigation and use by non-experienced users. All the data can be downloaded as standalone text files. ProtMiscuity will be updated on a regular basis as new evidence becomes available.

## Usage

ProtMiscuity can be searched by protein name or Uniprot ID, by organism or by the name of its canonical or promiscuous activities. An index of proteins is also available to browse. A typical query using the protein name retrieves general information about it in the form of browsable cards, including the protein family, source organism, the number of promiscuous and canonical reactions in which it is involved and the number of structures related. Searching with a molecule name or putative substrates/products of catalysis retrieve all proteins linked with the query or with similar molecules (Figure 1). By clicking on a protein the user is directed to its dedicated page, which displays detailed information on the protein, including its canonic and promiscuous reaction sites mapped (using ProViz (Jehl *et al.*, 2016)) onto sequences and known structures.

**Figure 1.**
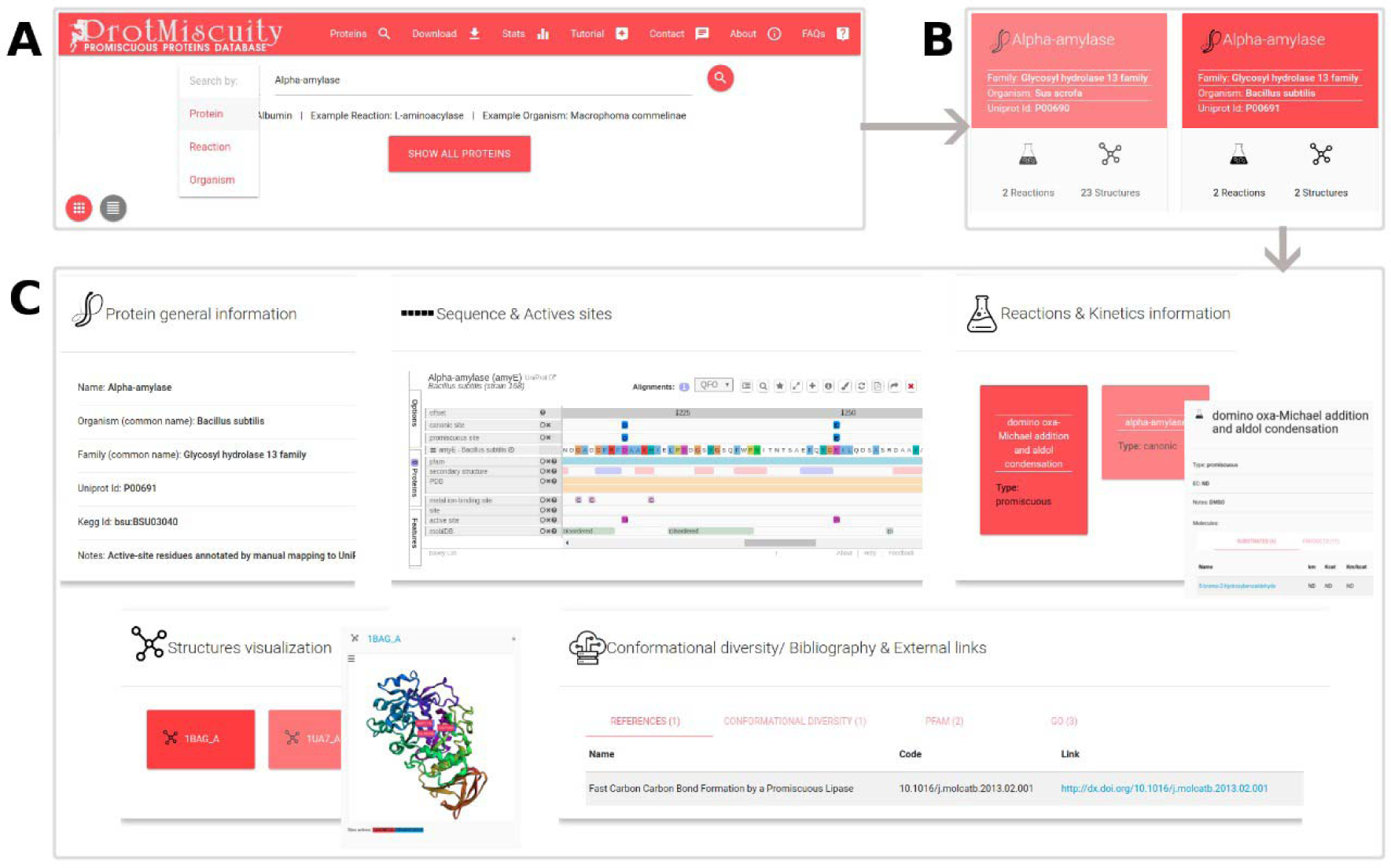
A) Home page of ProtMiscuity. The database can be searched using protein names, organism or target reaction. In this example, a search for the protein alpha-amylase is performed. B) Results page. It shows all matches to the query term in the form of protein-specific cards. In this example, alpha-amylases from two distinct organisms are retrieved. C) Information page. Clicking on one protein’s card displays to all available information for it, organized in five sections of interest. From top to bottom, left to right: a general description of the protein; the mapping of the canonic and promiscuous active sites, along with other source of relevant information, on the protein’s sequence; information about canonic and promiscuous activities, with known substrates, products and kinetic parameters (top panel); a visualization of each available structure of the protein, with catalytic sites mapped on it; and examples of conformational diversity, plus links to relevant bibliography and other databases, as separate tabs (bottom panel).

## Conclusions

Understanding the origin and mechanisms related with promiscuity may be a key feature for a deeper interpretation of protein function and evolution. Characterization of promiscuous behaviour has broaden the chemical repertory of enzymatic reactions, uncovering a large number of potential applications in biotechnology and related areas (López-Iglesias and Gotor-Fernández, 2015; Bornscheuer and Kazlauskas, 2004). ProtMiscuity provides a useful dataset for exploring new putative catalytic activities and their underlying mechanisms, as well as for retrieving complete information to develop and test new computational tools for the study and prediction of promiscuous behaviour.

## Acknowledgements

The authors would like to thank Dr. Norman Davey for his help with integrating Proviz into the ProtMiscuity web server and to both Giuliano Greco and Alexander Monzon for their help in designing and deploying the website.

